# quasitools: A Collection of Tools for Viral Quasispecies Analysis

**DOI:** 10.1101/733238

**Authors:** Eric Marinier, Eric Enns, Camy Tran, Matthew Fogel, Cole Peters, Ahmed Kidwai, Hezhao Ji, Gary Van Domselaar

## Abstract

**Summary:** quasitools is a collection of newly-developed, open-source tools for analyzing viral quasispcies data. The application suite includes tools with the ability to create consensus sequences, call nucleotide, codon, and amino acid variants, calculate the complexity of a quasispecies, and measure the genetic distance between two similar quasispecies. These tools may be run independently or in user-created workflows.

**Availability:** The quasitools suite is a freely available application licensed under the Apache License, Version 2.0. The source code, documentation, and file specifications are available at: https://phac-nml.github.io/quasitools/

## Introduction

The existing predominance and regular emergence of viral pathogens represent a significant public health threat worldwide. The ability to rapidly identify and characterize viral disease outbreaks is an essential component of public health response. Many effective conventional molecular methods exist and are routinely applied for viral disease detection, surveillance, and characterization. However, next-generation sequencing (NGS) technologies offer a far more sensitive, accurate, and comprehensive means for studying, characterizing, and monitoring viral pathogens.

Viral quasispecies are viruses that replicate in high numbers but with low fidelity, which results in a complex and dynamic spectrum of mutations within an infected individual [1]. Analysis of viral quasispecies poses a unique computational challenge, owing to the unprecedented data volume and complexity of viral quasispecies, sequence variations at diverse frequencies across the genome, demanding quality control strategies required for NGS data processing, and a lack of consensus in performing such analyses. Bioinformatics efforts to address the unique challenges of virology thus far have been underwhelming relative to their biomedical value and public health importance [2].

Nevertheless, there are a number of tools that exist for analyzing viral quasispecies. There are many tools that estimate the global diversity of viral quasispecies using read graph-based methods [3–7], probabalistic clustering methods [8–10], and *de novo* assembly methods [11]. HIVE [12] is a web environment that contains a tool for population analysis of viral quasispecies. The tool correlates distant mutations, identifies clones of low coverage, and outputs diagrams visualizing this information. PAQ [13] is capable of partitioning quasispecies sequences into groups that are genetically similar. Beyond these examples, there are numerous pipelines for variant calling and drug resistance identification [14–20]. Many applications designed for quasispecies analysis deal with specific issues in resolving viral quasispecies. The motivation behind quasitools is to provide a single collection of open-source, general-purpose tools for quasispecies analysis that are designed to operate seamlessly with each other.

## Implementation

quasitools is a Python application that provides several tools for quasispecies analysis, including variant calling, consensus generation, and reporting of viral quasispecies complexity (Table 1). These tools may be run independently or together in user-created workflows. Here we provide a brief summary of some of these tools.

**Table 1:** An overview of the tools available in quasitools.

| Tool | Description |
| --- | --- |
| aacoverage | builds an amino acid consensus and returns its coverage |
| call aavr | calls amino acid mutations for a BAM file and a supplied reference file |
| call codonvar | calls codon variants for a BAM file and a supplied reference file |
| call ntvar | calls nucleotide variants for a BAM file and a supplied reference file |
| complexity | reports the complexity of a quasispecies using several measures |
| consensus | generates a consensus sequence from a BAM file |
| distance | measures the distance between two quasispecies using angular cosine distance |
| dnds | calculates the dN/dS value for each region in a BED file |
| drmutations | identifies AA mutations corresponding to known drug resistance mutations |
| hydra | identifies HIV drug resistance mutations in an NGS dataset |
| quality | performs quality control on FASTQ reads |

The *quality* tool performs basic quality control on read sequencing data. It can filter FASTQ-formatted reads based on average quality score, median quality score, and read length. It can also mask low-quality bases and perform iterative read trimming. The *consensus* tool generates the consensus sequence of a quasispecies, using either a most-frequent base strategy or using a strategy that reports ambiguous IUPAC bases at positions where there is insufficient agreement.

There are multiple variant calling tools that can identify nucleotide (*call ntvar*), codon (*call codonvar*), and amino acid variants (*call aavar*) within a quasispecies, given a BAM alignment file and a FASTA reference file. The output of these variant calling tools may be provided to other tools in order to identify drug-resistant mutations.

We have developed a tool, *complexity*, which implements a wide variety of quasispecies complexity measurements for haplotypes, as described by Josep et al. 2016 [1]. These multidimensional measures capture the number of mutants, frequency of different haplotypes, and viral population size. Additionally, we have extended this framework for evaluating quasispecies complexity from amplicons to *k*-mers across a sequence pileup. This enhancement enables the study of quasispecies complexity across the genome using NGS data, but is limited by the length and accuracy of the sequencing reads.

We provide a tool, *distance*, for estimating the genetic distance between two quasispecies, which may be a useful and realtively quickly derived approximation for evolutionary relatedness among quasispecies of the same species. The tool represents each quasispecies pileup as a vector and calculates the angular cosine distance between each of these vectors. We use angular cosine distance as it is accommodates vectors that are both sparse and varied in their depth of coverage. Furthermore, the user may normalize the pileup vectors to prevent bias in relative coverage within a single pileup, where a region with exceptional coverage would dominate the distance calculation.

HyDRA is an annotated, reference-based pipeline, which analyzes NGS data for genotyping HIV-1 drug resistance mutations. HyDRA combines the analyses of many tools in quasitools in order to identify drug resistance mutations in HIV quasispecies. It uses an annotated HXB2 sequence for reference mapping and stringent variant calling to identify HIV drug resistant mutations based on the Stanford HIV Drug Resistance Database and the 2009 WHO list for Surveillance of Transmitted HIV Drug Resistance. As HyDRA is built using the tools available within quasitools, it is possible for users to use quasitools to create similar workflows for other viral quasispecies.

The quasitools package is freely available as source code, as a Conda package, and individually as tools on the public Galaxy Tool Shed for use on the Galaxy web-based platform.

## Funding

This work was funded by a grant from the Canadian Federal Government Genomics Research and Development Initiative (GRDI).

## Conflict of Interest

None declared.

